# On triangular Inequalities of correlation-based distances for gene expression profiles

**DOI:** 10.1101/582106

**Authors:** Jiaxing Chen, Yen Kaow Ng, Lu Lin, Yiqi Jiang, Shuaicheng Li

## Abstract

Various distance functions for evaluating the differences between gene expression profiles have been proposed in the past. Such a function would output a low value if the profiles are strongly correlated—either negatively or positively—and vice versa. One popular distance function is the absolute correlation distance, *d*_*a*_ = 1 − *|ρ|*, where *ρ* is some similarity measures, such as Pearson or Spearman correlation. However, absolute correlation distance fails to fulfill the triangular inequality, which would have guaranteed better performance at vector quantization, allowed fast data localization, as well as sped up data clustering. In this work, we propose 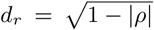 as an alternative. We prove that *d*_*r*_ satisfies the triangular equality when *ρ* represents Pearson correlation, Spearman correlation, or Cosine similarity. We empirically compared *d*_*r*_ with *d*_*a*_ in gene clustering and sample clustering experiment, using real biological data. The two distances performed similarly in both gene cluster and sample cluster in hierarchical cluster and PAM cluster. However, *d*_*r*_ demonstrated more robust clustering. According to bootstrap experiment, the number of times where *d*_*r*_ generated more robust sample pair partition is significantly (p-value *<* 0.05) larger. This advantage in robustness is also supported by the class “dissolved” event.

## 1 Introduction

In biological data analysis we are frequently required to evaluate how similar two genetic expression profiles are. For example, when identifying gene expression patterns across different conditions, when clustering genes of similar functions [6, 10], when detecting the gene temporal profile of relevant functional categories by time-series data clustering [8], when measuring similarity between genes in microbial community [5], and when inferring gene regulatory network [20].

Several distance functions are currently used to evaluate this similarity—the most prominent one being the absolute correlation distance. The function regards positive correlation and negative correlation equally, giving a value of zero to highly correlated profiles (whether positively or negatively correlated), and a value of one to uncorrelated profiles. More precisely, the absolute correlation distance is defined as *d*_*a*_= 1−*|ρ|*, where *ρ* can be Pearson correlation, Spearman correlation, uncentered Pearson correlation (which is equivalent to Cosine similarity), or Kendell’s correlation. Profiles which are highly correlated have *ρ* = 1 or *ρ* = −1, and hence resulting in *d*_*a*_= 0; profiles which are unrelated have *ρ* = 0, hence resulting in *d*_*a*_= 1. The absolute correlation distance is widely used, for example, in measuring the co-expression similarity between the profiles of genes in WGCNA [18], clustering of gene expressions [4], and in defining the abundance similarity between OTUs in microbiome area [5]. However, in spite of its widespread usage, it has been noted that most variants of the measure, with the exception of the absolute Kendall’s correlation, suffer from the drawback of not satisfying the triangular inequality [8, 12].

A distance measure *d* which (1) satisfies the triangular inequality and (2) has *d*(*x, y*) = 0 when *x* = *y*, is called a *metric* [1, 21]. Researchers have observed that the performance of vector quantization improves when the measure used is a metric [23]. A measure which fulfills triangular inequality would allow faster data localization as well as speed up data clustering [1, 22]. Many clustering algorithms, such as k-means [7] and DBSCAN [17], can exploit triangular inequality to achieve better performance. For instance, a distance calculation can be skipped as soon as it is found to exceed lower or upper bounds estimated through triangular inequality [7]. The same strategies cannot be applied on distance measures that violate triangular inequality without compromising the quality of the clustering [1].

Variants of the absolute correlation distance are not the only distance measure used in gene expression analysis that violate triangular inequality. Prasad *et al.* [24] compiled a list of distance measures for analysis on gene expression profiles. Many of the measures in the list do not fulfill triangular inequality. These include the Harmonically summed euclidean distance, Bray-Curtis distance, Pearson correlation distance, absolute Pearson correlation distance, uncentered correlation distance, absolute uncentered correlation distance, Pearson linear dissimilarity, Spearman correlation distance, absolute Spearman rank correlation, and the Cosine distance.

In this work, we propose an alternative *d*_*r*_ to the absolute correlation distance, defined as 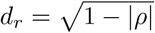, where *ρ* can be Pearson correlations, Spearman correlations, or uncentered Pearson correlation (or Cosine similarity). We show that *d*_*r*_, unlike *d*_*a*_, satisfies the triangular equality for all of these correlations.

We compared the performance of *d*_*r*_ to *d*_*a*_ in biological data clustering. The clustering method includes hierarchical clustering and PAM (partitioning around medoids) [16]. For *ρ* we used Pearson correlation, Spearman correlation, and Cosine similarity. As data we used 16 normalized time-series datasets and cancer samples cluster in 35 expression datasets. Performances for the sample cluster tests were evaluated with adjusted Rand index (ARI) [25], while those for the gene cluster tests were evaluated with functional analysis.

Our result shows the two distance measures led to identical hierarchical cluster partition in complete linkage and single linkage, but different in average linkage. In the gene cluster experiment, *d*_*r*_ outperformed *d*_*a*_ in 10, 9, and 10 datasets among 16 datasets for average linkage hierarchical cluster, and 9, 12, 7 for PAM experiment. In sample cluster experiment, *d*_*a*_ and *d*_*r*_ obtained the same ARI in at least 27 datasets among all 35 sample cluster dataset. The two distances have comparable performances in real gene cluster and sample cluster, although the clustering performed with *d*_*r*_ are more robust than those with *d*_*a*_. When tested with multiple bootstrap test, *d*_*r*_ outperformed *d*_*a*_ at robustness. *d*_*r*_ led to more robust clusters than *d*_*a*_ in both hierarchical cluster, when considering internal nodes, and PAM cluster when any of the correlations is used as *ρ*. For PAM clustering with Pearson correlation used as *ρ*, in more than 34 datasets, *d*_*r*_ generated significantly (p-vlaue *<* 0.05) more robust sample pair partition than *d*_*a*_. Similar results were obtained when *ρ* is Spearman correlation and Cosine similarity. The robustness of *d*_*r*_ is also supported by statistics on the time a class “dissolved”.

We also compare *d*_*r*_ to other variants of *d*_*a*_ where *ρ* is squared [27], that is,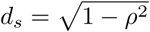. Our results showed *d*_*r*_ to have better performance at clustering.

## 2 Method

### 2.1 Prove triangular inequality of the transformation on Pearson correlation

The original absolute correlation distance *d*_*a*_= 1 *− |ρ|* dissatisfy triangular inequality. We propose a new measure *d*_*r*_, as

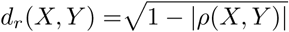

where *X* and *Y* are expression profiles, and *ρ* can be any one of Pearson correlation coefficient, Spearman correlation, or uncentered Pearson correlation.

We first show that *d*_*r*_ is a metric. Take X=(*x*_1_, *x*_2_, *…, x*_*n*_), Y=(*y*_1_, *y*_2_, *…, y*_*n*_) and Z=(*z*_1_, *z*_2_, *…, z*_*n*_), then the triangular inequality can be written as

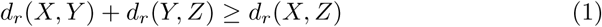

This demonstrates that *d*_*r*_ satisfies the triangular inequality, when *ρ* is Pearson correlation coefficient, Spearman correlation or the uncentered Pearson correlation, according to the details in the supplementary material.

### 2.2 Evaluation

To compare our modified absolute Pearson correlation distance 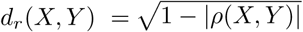 to the original absolute Pearson correlation distance *d*_*a*_(*X, Y*) =1–*|ρ*(*X, Y*) *|*, we performed clustering experiment on real microarray datasets, including 16 gene time-series profile datasets [15] and 35 datasets for clustering of cancer samples [26]. The clustering algorithms for the test include hierarchical clustering and PAM. The input of a clustering task is a distance matrix and the output is a partitioning which gives the clusters. We performed clustering by both gene and sample.

For the sample clusters, we selected the number of clusters, *k*, according to benchmark. We evaluated the clustering result by examining how consistent the clusters are with the benchmark by ARI [25]. A greater ARI value indicates higher concordance between the cluster partition and the benchmark partition. Given a partition *u* and a reference partition *v*,

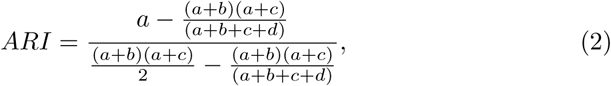

where *a* refers to the total number of sample pairs belonging to the same cluster in both *u* and *v, b* refers to the total number of sample pairs in the same cluster in *u* but in different clusters in *v, c* is the total number of sample pairs that are in different clusters in *u* but in the same clusters in *v*, and *d* refers to the total number of sample pairs that are in different clusters in both *u* and *v*.

For the gene clusters, we evaluated clustering performance by gene functional analysis [14]. The number of clusters was determined according to Calinski-Harabasz Index (*CH*_*index*_) [2] as follows. The *CH*_*index*_ is given as

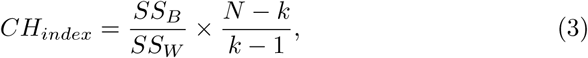

where *k* is the number of clusters, and *N* is the total number of samples, *SS*_*W*_ is the overall within-cluster variance, *SS*_*B*_ is the overall between-cluster variance. A higher *CH*_*index*_ value implies a better solution. We used the value of *k* which corresponds to the peak or at least an abrupt elbow on the line-plot of *CH*_*index*_ value.

After obtaining the clusters, we performed GO enrichment for each generated cluster with R package [3, 9, 13]. For each cluster generated by *d*_*a*_, we got a set of significant GO terms with p-value *<* 0.05, denoted as r1. Similarly, for cluster generated by *d*_*r*_, we got a set of significant GO term r2. After that, for two result list r1 and r2, we counted the number of times that the GO term of r1 has smaller p-value than that of r2, denoted as ≠ (*r*1 *< r*2), and the number of times that GO term of r2 has smaller p-value than it of r1, denoted as ≠ (*r*2 *< r*1). Then we calculated

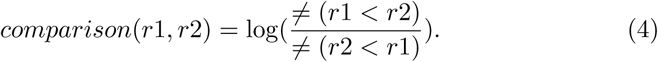

Positive values of *comparison*(*r*1, *r*2) imply that r1 is better than r2, and negative values imply the opposite. So the negative values mean *d*_*r*_ wins *d*_*a*_ in this dataset. If we change the order of the results under comparison (r1,r2) or (r2,r1), it will only change the sign of the result, but not its absolute value.

### 2.3 Robustness test

To test the robustness of cluster with different distance measures, we performed bootstrap experiments on the 35 microarray datasets in clustering cancer samples, and investigated the “dissolved” [11] event for class given by different cluster processes. For each dataset, we first obtained an original partition *p*_*o*_ from the original dataset. Then, for each dataset, suppose there are *n* samples in total, we bootstraped 100 times. For each time, we randomly selected *n* samples with replicate from the original dataset, and performed clustering on the resampled data to get a resulting partition *p*_*i*_. We compared *p*_*i*_ with *p*_*o*_. Denote *c*_*oj*_ as class *j* in *p*_*o*_, *c*_*ik*_ as class *k* in *p*_*i*_. For each *i*, we calculated the Jaccard similarity, *J*_*ijk*_, between each *c*_*oj*_ and all *c*_*ik*_. Then we calculated *J*_*ij*_= max(*J*_*ijk*_). After repeating 100 times, for each *c*_*oj*_ in *p*_*o*_, we obtained 100 similarity values, respectively denoted *J*_*ij*_ for each of the bootstraps. If *J*_*ij*_ *<* 0.5, we take *c*_*oj*_ as having “dissolved” in bootstrap *i*. We counted the number of times the class *c*_*oj*_ dissolved in 100 bootstrap. If this frequency is larger than 40, we regard *c*_*oj*_ as being dissolved in the experiment. We repeated the bootstrap process for multiple iterations. We tested the robustness of the class by comparing the times it dissolved in multiple iterations. Finally, we compared the performance of *d*_*a*_ and *d*_*r*_ by comparing the robustness of the classes they generated.

We also investigated sample pairs for robustness. We selected sample pairs that are clustered together to see whether they are consistently clustered together across multiple runs, in which case, the result for the sample pair is robust. Similarly we examined sample pairs that are not clustered together to see if they are consistently placed in different classes. For each sample pair *i* and *j* in one dataset, we counted the number of times *n*_1_ they are sampled together in 100 bootstraps, the number of times *n*_2_ they are clustered in the same class, and the number of times *n*_3_ they are clustered in different classes. If *n*_2_ *> n*_3_, then this pair is decided as consistently clustered, otherwise they are not consistently clustered. For each sample pair, we calculated the ratios *n*_2_*/n*_1_ as well as the median value *m*_*together*_ for all the non-zero ratio values. This is repeated for *n*_3_*/n*_1_ and their median, *m*_*notTogether*_. Then we calculated *υ* = *m*_*together*_ *∗m*_*notTogether*_. A larger *υ* implies a more robust clustering. We recorded this as a “win” event for *d*_*r*_ if *υ*_*r*_ *> υ*_*a*_. For 35 files, we got a list of *υ* for *d*_*r*_ and *d*_*a*_. We did Wilcox test for the list of *υ*_*r*_ and *υ*_*a*_ with alternative hypothesis as true location shift is not equal to 0.To see wether *υ* for *d*_*r*_ is significantly larger than *υ* for *d*_*a*_.

## 3 Results

### 3.1 Performance in gene cluster

First, we evaluated the performance of 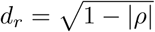and the *d*_*a*_= 1 −*|ρ|* on gene clustering. As data we used 16 time-series profile datasets with normalization from a previous work [15]. For each dataset we calculated the distances *d*_*r*_ and *d*_*a*_ for pairwise gene profiles, resulting in the distance matrices *M*_*r*_ and *M*_*a*_. Then, we applied hierarchical clustering and PAM for each distance matrix, estimating the number of cluster *k* by *CH*_*index*_. Since the data sets do not have a reference partition for genes, we evaluated the performance with biological functional analysis [14]. The clustering result with a higher scored GO term is considered as the better solution. For the hierarchical clusters, we tested three modes, namely complete linkage, single linkage, and average linkage. Clustering using either *d*_*r*_ or *d*_*a*_ led to identical dendrograms in complete linkage hierarchical clustering as well as in single linkage.

For hierarchical clustering with average linkage, *d*_*r*_ outperformed *d*_*a*_ in 10, 9, and 10 datasets among 16 datasets when *ρ* is any of Pearson correlation, Spearman correlation, and Cosine similarity respectively (see Fig. 1). In PAM experiments, *d*_*r*_ outperformed *d*_*a*_ in 9, 12, 7. The two measures outperformed each other for nearly equal number of times.

**Fig 1.**
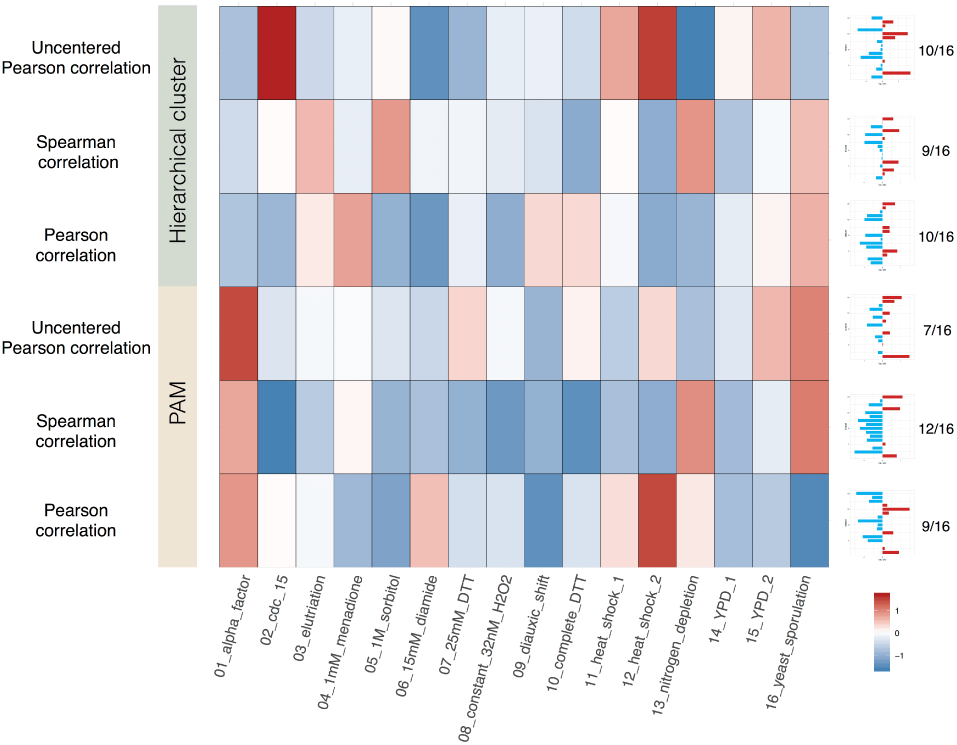
Result of comparing *d*_*r*_ and *d*_*a*_ in gene clustering. Each column corresponds to one time-series profile dataset. Each row corresponds to one comparison between *d*_*r*_ and *d*_*a*_ while *ρ* is different correlation in certain clustering method. Color refers to the value of *comparison*(*r*1, *r*2). Negative value implies that *d*_*r*_ is better than *d*_*a*_, and positive values implies the opposite. For each comparison combination, there is a barplot on the right side on the corresponding row. The x-axis of the barplot refers to *comparison*(*r*1, *r*2).

### 3.2 Performance in sample cluster

To compare the performance of *d*_*a*_ and *d*_*r*_ in sample clusters, we used 35 datasets from a previous work [26]. The samples in each dataset is assigned a label such as disease or healthy. We applied normalization to each dataset by scaling each gene to the standard normal distribution. We then performed hierarchical clustering and PAM, with the number of clusters *k* set as the number of the unique labels in each dataset. We evaluated the performance by ARI [25], which measures the consistency between cluster partition and benchmark labels.

For hierarchical cluster, the complete and single mode resulted in identical dendrograms. When *ρ* is Pearson correlation, the hierarchical cluster in average mode using both *d*_*a*_ and *d*_*r*_ resulted in similar ARI across all 31 datasets (see Fig. 2). When *ρ* is Spearman correlation or uncentered Pearson correlation, both *d*_*a*_ and *d*_*r*_ resulted in the same ARI in at least 27 datasets, for both methods of clustering. For those datasets with different ARI in comparison, the number of times *d*_*r*_ outperforms *d*_*a*_ is close to the number of times *d*_*a*_ outperforms *d*_*r*_. As an example, in PAM *d*_*r*_ outperformed *d*_*a*_ 5 times while *d*_*a*_ outperformed *d*_*r*_ 3 times when *ρ* is Spearman correlation. These results show that they have comparably good performance in our sample cluster experiment.

**Fig 2.**
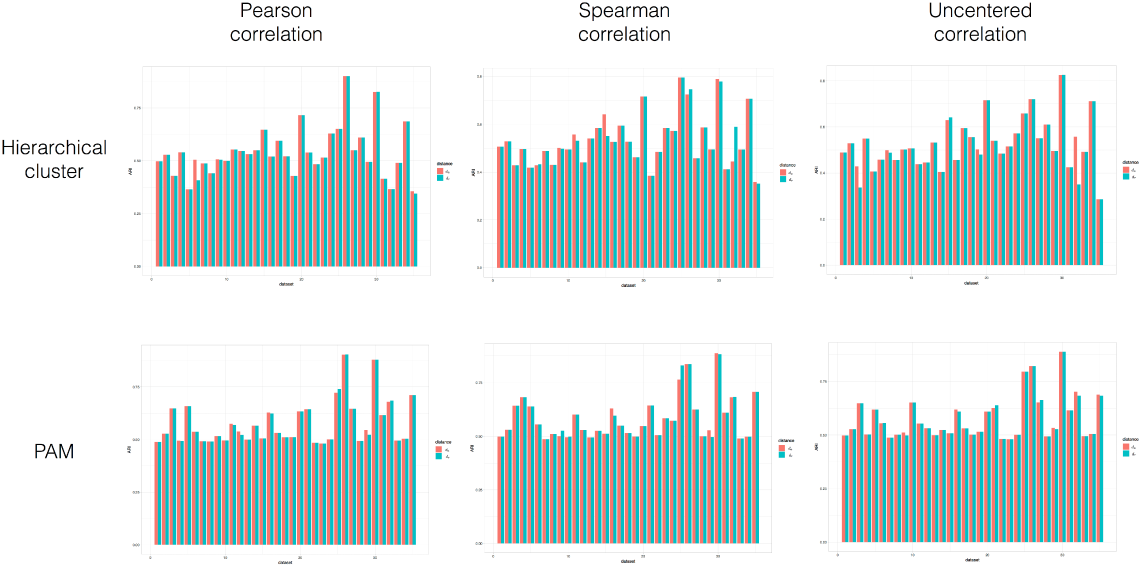
Result of comparing *d*_*r*_ and *d*_*a*_ in sample clustering. For each subfigure, x-axis refers to different datasets, y-axis refers to the ARI value. A larger ARI implies a better partitioning.

### 3.3 Detailed analysis with an example

To see how *d*_*r*_ and *d*_*a*_ lead to different cluster results, we used one dataset as an example and observed the clustering process. We used the 18-th dataset [19] in the sample cluster experiment, performing hierarchical clustering, using Pearson correlation as *ρ*.

In the beginning, the distance matrices *M*_*r*_ and *M*_*a*_ calculated according to *d*_*r*_ and *d*_*a*_ are the same in rank, in the sense that if we sort the values in *M*_*r*_ and *M*_*a*_ increasingly, the two lists will have the same order. For hierarchical clustering with the complete linkage and single linkage, *d*_*r*_ and *d*_*a*_ led to the same resultant dendrogram because they only take maximum or minimum distance value when calculating the distance between cluster, thus introducing no new value of distance during the entire clustering process. For hierarchical cluster with average linkage, the same two samples are merged at the first step, thus the smallest distances are combined into one cluster. Since an average distance is computed of the newly generated cluster, a difference in rank emerges. In Fig. 3A, the circle network shows the pairs which are different in the ranks of the distance sets generated by *d*_*a*_ and *d*_*r*_ in step 2 to step 6. In this dataset, *d*_*a*_ and *d*_*r*_ led to the same ARI even though the resultant dendrograms are different in structure. The dendrogram for *d*_*a*_ is shown in Fig. 3B and that for *d*_*r*_ in Fig. 3C. The difference between the two dendrograms is colored in red. Fig. 3D shows the distribution of the ranks which are different.

**Fig 3.**
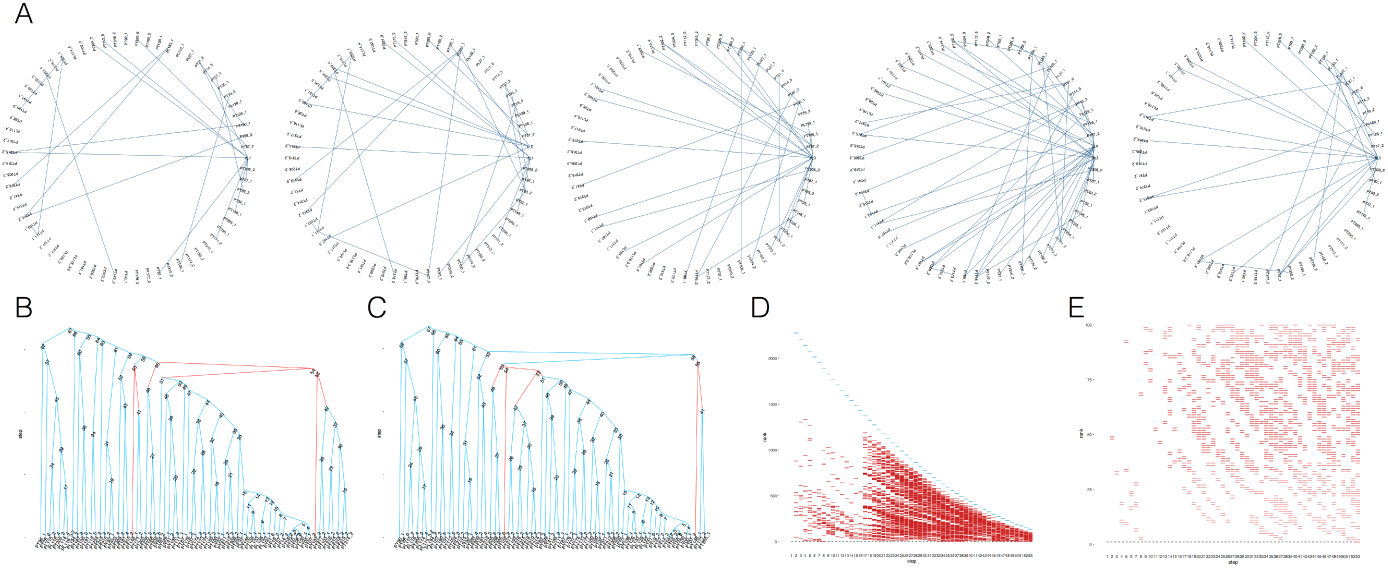
An example for average linkage hierarchical cluster with *d*_*a*_ and *d*_*r*_. A. The circle networks show the pairs where the distance is different in the ranks generated by *d*_*a*_ and *d*_*r*_ in step 2 to step 6. Nodes in network refer to samples for clustering. Edges refer to the distance where two sample are different in rank in *d*_*a*_ and *d*_*r*_. *c*1 refers to the the class generated in step 1, *c*_2_ refers to the class generated in step 2. B. Dendrogram for *d*_*a*_. C. Dendrogram for *d*_*r*_. The difference between the two dendrograms is colored in red. D. Distribution of ranks which are different in *d*_*a*_ and *d*_*r*_. E. Zoom in for the top 100 rank for Figure 3D.

From step 2 to step 52, there exist different ranks in the distance of pairs in two distance experiments. However, the pairs of rank 1 are the same, showing that both *d*_*a*_ and *d*_*r*_ led to the same two samples being merged into a new cluster. In step 53, the pair of rank 1 started to differ, showing that different samples in two distance experiments have been selected. This difference is reflected in the resultant dendrogram. As shown in Fig. 3B and Fig. 3C, for *d*_*a*_, *c*_42_ and “*PT* 102_2_” have been merged, while for *d*_*r*_, *c*_42_ and *c*_51_ have been merged (*c* represents the internal node in the dendrogram).

In the sample cluster experiment, due to scarcity in the number of pairs (the maximum number of samples in a single dataset is 248 among datasets in this sample cluster experiments), the difference in ARI only occurred in 4 out of 35 datasets. In gene cluster experiment, the boosted number of pairs enlarged the differences in the dendrogram, hence the partition is different in all 16 time-series datasets.

### 3.4 Robustness test

We compared the methods’ robustness with bootstrap experiments in clustering cancer samples on 35 microarray datasets. This is done by examining the number of sample pairs that are consistently clustered across 20 iterations. In each iteration, we resampled 100 times for each dataset. For PAM, *d*_*r*_ displayed more robust clustering than *d*_*a*_. Fig. 4A, B, C and D are for comparing *d*_*r*_ and *d*_*a*_ through PAM clustering using Pearson correlation as *ρ*. Fig. 4A shows the number of times *d*_*r*_ achieved a win over 20 iterations in each dataset. *d*_*r*_ achieved more win in 34 datasets among 35 datasets (see Fig. 4B). Fig. 4C shows the box plot for *υ* over 20 iterations and 35 datasets.

**Fig 4.**
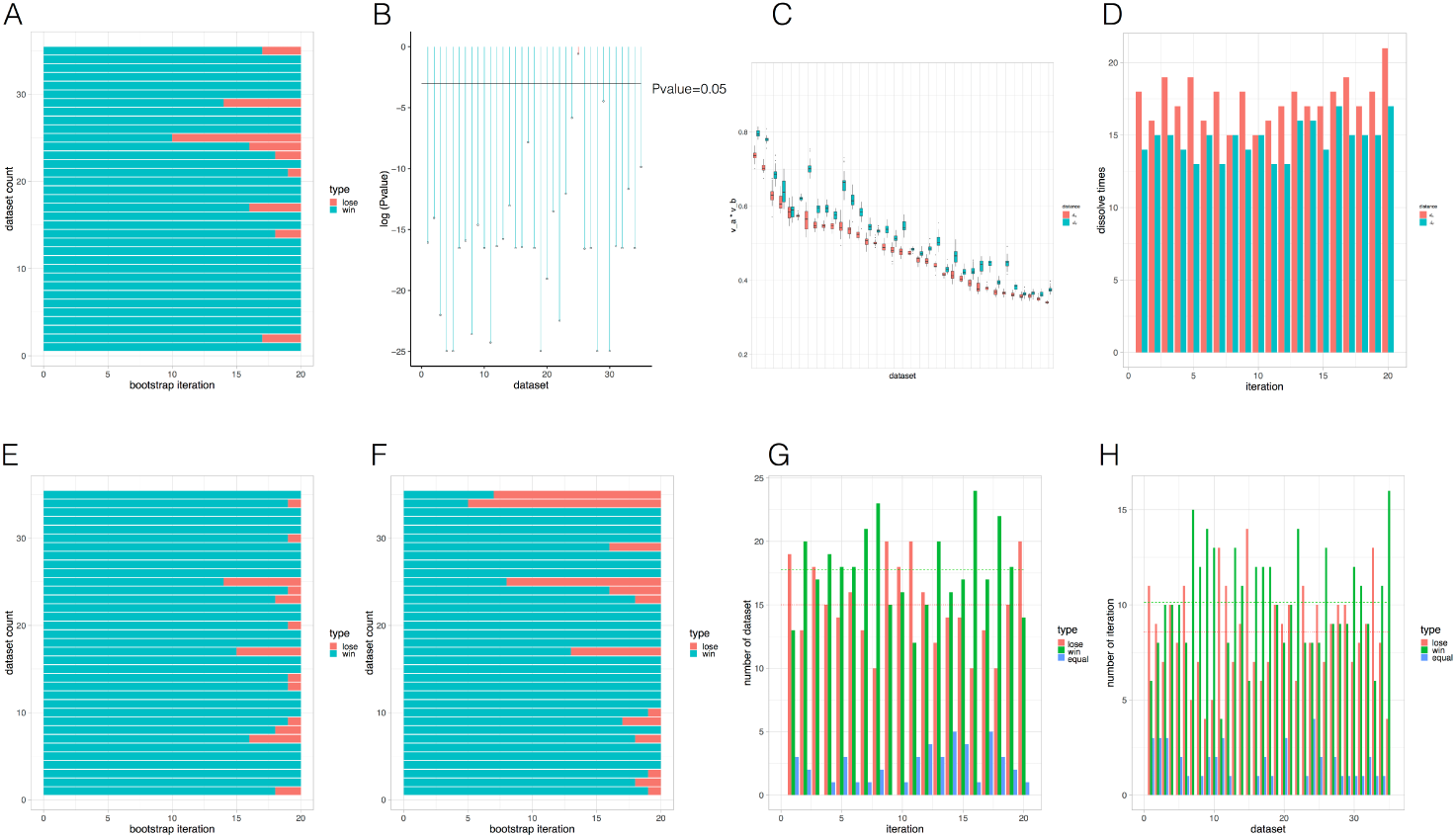
Result for robustness test on *d*_*a*_ and *d*_*r*_. A,B,C,D. Results obtained using Pearson correlation as *ρ* on PAM. A. The number of times *d*_*r*_ win over 20 iterations in each dataset. Each row corresponds to one dataset. B. p-values in testing the difference between the number of times *d*_*r*_ wins in all 35 datasets. Each point corresponds to one dataset. C. Each box represents one *υ* value over 20 iterations per dataset. We compared the box plot for *d*_*a*_ and *d*_*r*_ in each dataset. The datasets in C have been reordered to fit the decrease of y value to show the trend more clearly. D. The number of classes “dissolved” in *d*_*a*_ and *d*_*r*_ across all 20 iterations. E. Result for Spearman correlation as *ρ* in PAM clustering. F. Result for Uncentered Pearson correlation as *ρ* in PAM clustering. G, H Results for Pearson correlation as *ρ* in hierarchical clustering, considering all internal nodes as classes, G. Result for comparing *d*_*a*_ and *d*_*r*_ by the number of times classes “dissolved” in 35 datasets over 20 iterations. The number of times *d*_*r*_ win, lose, or is equal to *d*_*a*_. The green horizontal line represents the average number across all the iterations where *d*_*r*_ wins. The red horizontal line represent the average number across all the iterations where *d*_*r*_ lose. H. Result for comparing *d*_*a*_ and *d*_*r*_ per dataset.

Fig. 4D shows the results where we evaluated robustness through the number of times a class is “dissolved”. The number of classes dissolved through *d*_*a*_ is larger than it in *d*_*r*_ in all 20 iterations. Hence, *d*_*r*_ led to more robust clustering results, consistent with our earlier results in Fig. 4A, B, C. Similar results are obtained when *ρ* is Spearman correlation and Cosine similarity, as shown in Fig. 4E and Fig. 4F.

For the hierarchical cluster, we examined all the internal nodes for the number of times those class dissolved for each dataset. Fig. 4G shows the number of datasets where *d*_*r*_ achieved a win. Fig. 4H shows the comparison according to each dataset. Both figures show that *d*_*r*_ achieved a win for more times than *d*_*a*_. Across 20 iterations, the average number of times when *d*_*r*_ wins is larger than the time *d*_*a*_ wins. In summary, the use of *d*_*r*_ resulted in more robust clustering than *d*_*a*_ in both hierarchical and PAM clustering.

## 4 Discussion

Failure in satisfying the triangular inequality is a severe problem in absolute correlation distance. We show how frequently this violation occurs in the 35 sample cluster datasets [26] (see Fig.S1 in supplementary material). The distributions differ across datasets, with fewer violations after normalization. The number of violations also appear to decrease during the merge process in average linkage hierarchical cluster. More violations (of up to 40%) appeared in the 16 gene cluster dataset, as shown in Fig.S1D.

Besides, we also compare *d*_*r*_ to squared correlation distance. In [27], two variants of the absolute correlation were proposed, namely 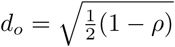 and 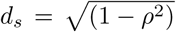, where Pearson correlations is used as *ρ*. These efforts would result in metric distances. The first variant, *d*_*o*_, has a range of 0 to 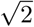, which results in inconsistencies with *d*_*a*_, thus limiting its use. The squared correlation distance, *d*_*s*_, on the other hand, is analytically less sensitive than *d*_*r*_ in responding to changes in *ρ*. This observation is confirmed by our empirical tests using hierarchical clustering (see Fig.S2 in supplementary material). In the tests, *d*_*r*_-based clustering outperformed *d*_*s*_ in 15 datasets, while losing out to *d*_*s*_ in only 8.

## 5 Conclusion

The absolute correlation distance *d*_*a*_=1–*|ρ|* is widely used in biological data clustering in spite of its shortcoming of not satisfying the triangular inequality. In this paper we proposed an alternative, *d*_*r*_, that does. Our comparison of *d*_*r*_ and *d*_*a*_ on gene clustering using 16 normalized time-series datasets and sample cluster in 35 expression datasets shows that the two distance measures led to identical clusters in hierarchical clustering with complete linkage and single linkage. The two distances have comparable performances in both gene cluster and sample cluster, using both hierarchical as well as PAM cluster, although *d*_*r*_-based clustering led to more robust clustering. The robustness of *d*_*r*_-based clustering is also supported by evaluation based on the number of times that a class “dissolved”.

## Supplementary material

### Supplementary figures

**Fig S1.**
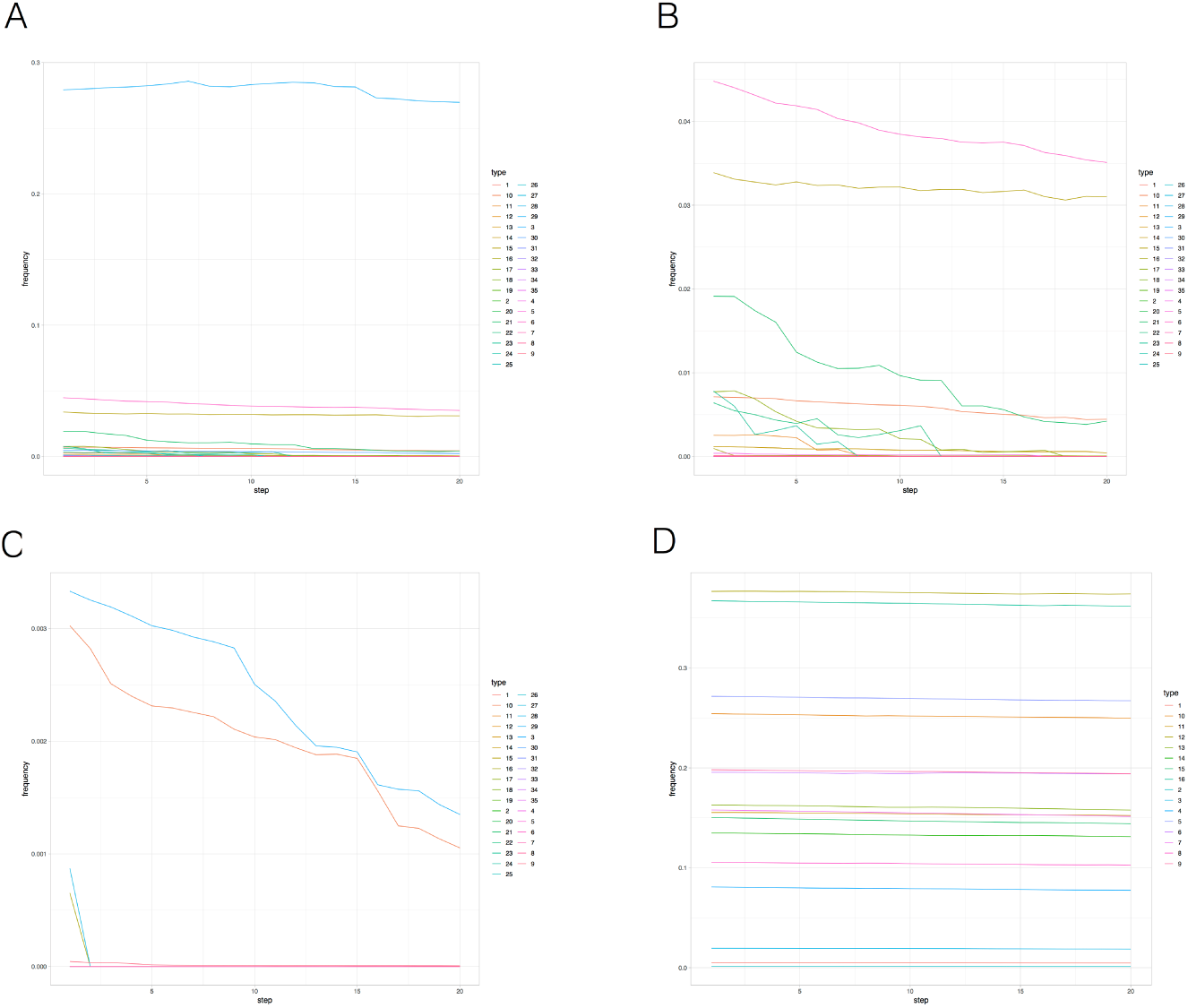
Percentage of pairs which dissatisfies triangular inequality under *d*_*a*_. If the distance between any pair of points fails to observe the triangular inequality for some third point, we consider the pair to have failed triangular inequality; the percentage of pairs that do not satisfy triangular inequality is shown. A. Percentage of pairs which dissatisfies triangular inequality, from step 2 to step 20 in hierarchical clustering, within the 35 sample cluster datasets without normalization. B. Detailed view of A. C. Percentage of pairs for the dataset with normalization, with scaling, for each gene. D. Percentage of pairs in hierarchical clustering in gene clustering.

**Fig S2.**
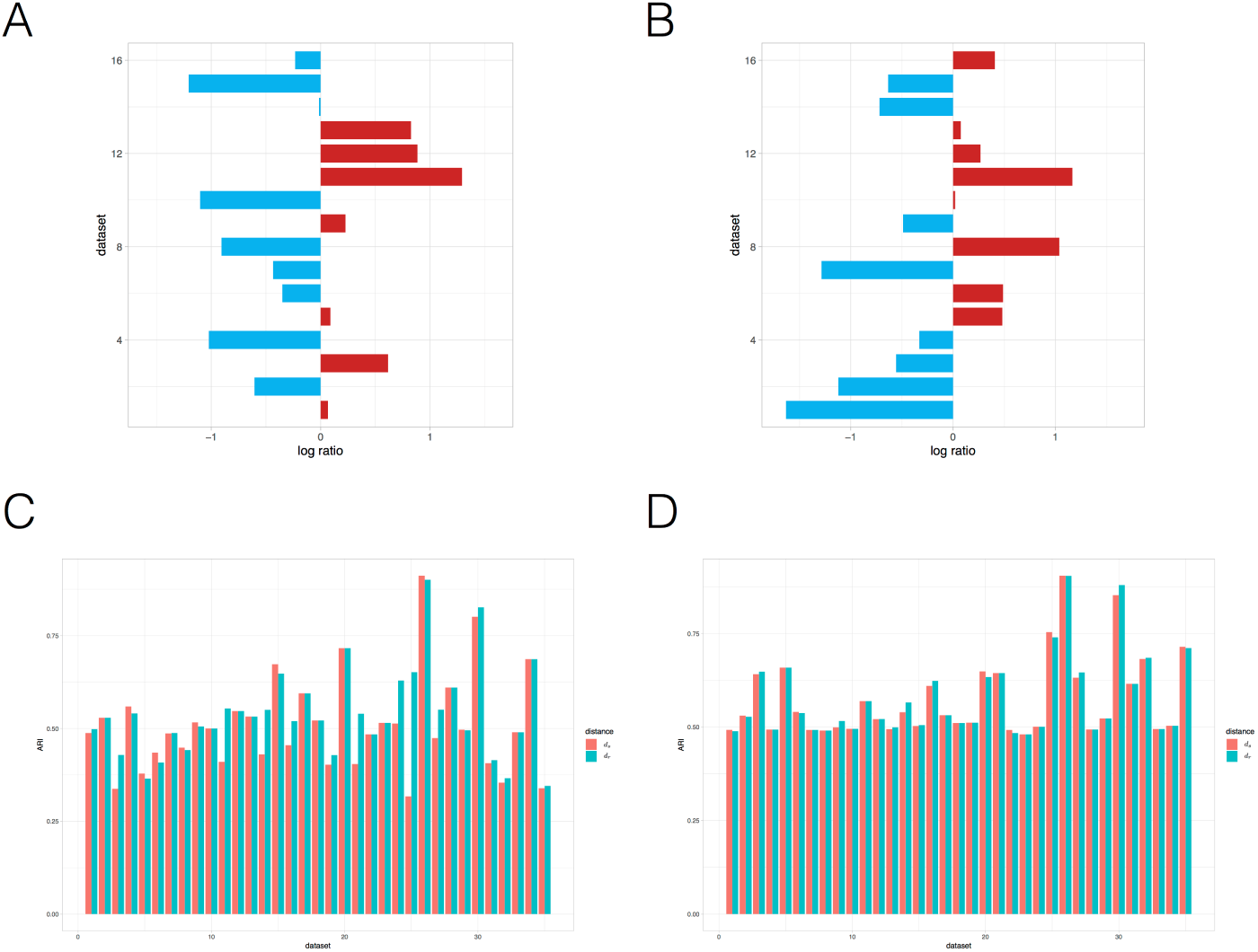
Result for comparing *d*_*r*_ and *d*_*s*_. Pearson correlation is used as *ρ*. A. Comparison on gene clustering using hierarchical cluster. X-axis refers to the value of *comparison*(*r*1, *r*2). Negative value implies that *d*_*r*_ is better than *d*_*a*_, while positive value implies that *d*_*s*_ is better. B. Comparison on gene clustering using PAM. C. Comparison on sample clustering using hierarchical cluster. Y-axis refers to ARI. D. Comparison on sample clustering using PAM.

### Proof of *d*_*r*_ fulfilling the triangular inequality for Pearson correlation as *ρ*

We define the distance of *X* and *Y* by 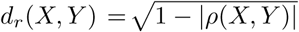, where *ρ* is the Pearson correlation coefficient.

By the triangular inequality of distance in n-dimensional Euclidean space, *d*_*r*_(*X, Y*) + *d*_*r*_(*Y, Z*) *≥ d*_*r*_(*X, Z*). Take *X* = (*x*_1_, *x*_2_, *…, x*_*n*_), *Y* = (*y*_1_, *y*_2_, *…, y*_*n*_) and *Z* = *z*_1_, *z*_2_, *…, z*_*n*_) such that

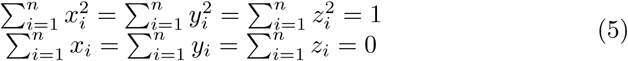

Then the triangular inequality

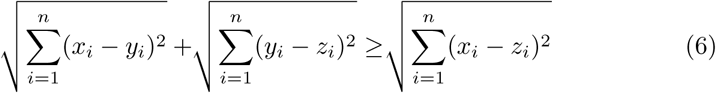

can be rewritten as

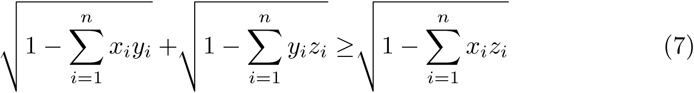

For data from a sample, the Pearson correlation coefficient can be calculated as follows

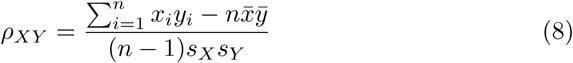

Since Pearson correlation coefficient is invariant under linear transformation, which means 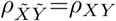with 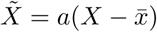 and 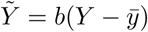 satisfying

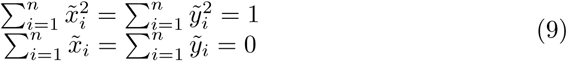

where 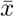 and 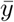 are the sample means of X and Y, it can be rewritten as

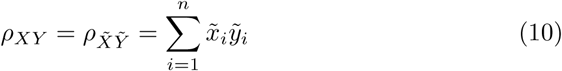

Without loss of generality, we assume that the samples are normalized (i.e. satisfying Equation 5).

Therefore, we have the modified Pearson distance

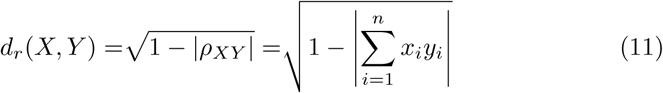

To prove the triangular inequality of *d*_*r*_, we divide this into eight cases by the signs of *ρ*.

**Case I** When *ρ*_*XY*_ *≥* 0, *ρ*_*YZ*_ *≥* 0, *ρ*_*XZ*_ *≥* 0,

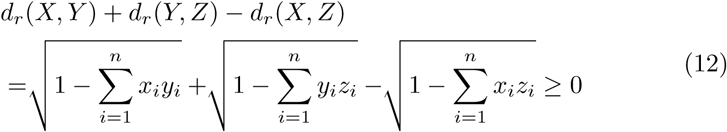

by (7)

**Case II** When *ρ*_*XY*_ *≥* 0, *ρ*_*YZ*_ *<* 0, *ρ*_*XZ*_ *<* 0, take *c*_*i*_= *-z*_*i*_.

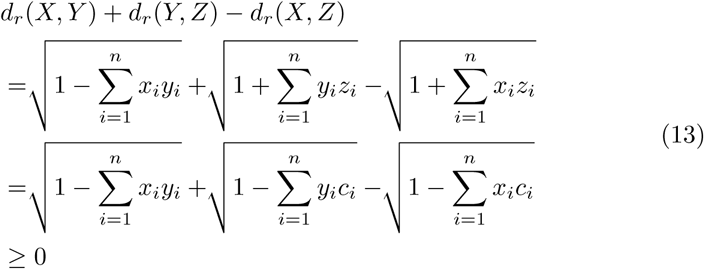

by (7)

**Case III** The case when *ρ*_*XY*_ *<* 0, *ρ*_*YZ*_ *≥* 0, *ρ*_*XZ*_ *<* 0 is equivalent to case II.

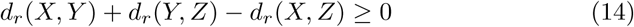

holds

**Case IV** When *ρ*_*XY*_ *<* 0, *ρ*_*Y*__*Z*_ *<* 0, *ρ*_*XZ*_ *≥* 0, take *b*_*i*_= *-y*_*i*_.

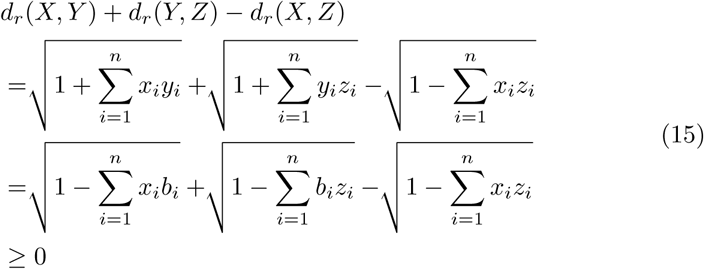

by (7)

**Case V** When *ρ*_*XY*_ *<*0, *ρ*_*Y*__*Z*_ *<*0, *ρ*_*XZ*_ *<*0

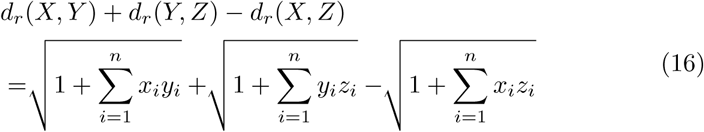

Take *b*_*i*_= *-y*_*i*_. Therefore we have

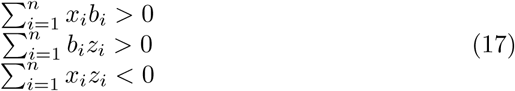

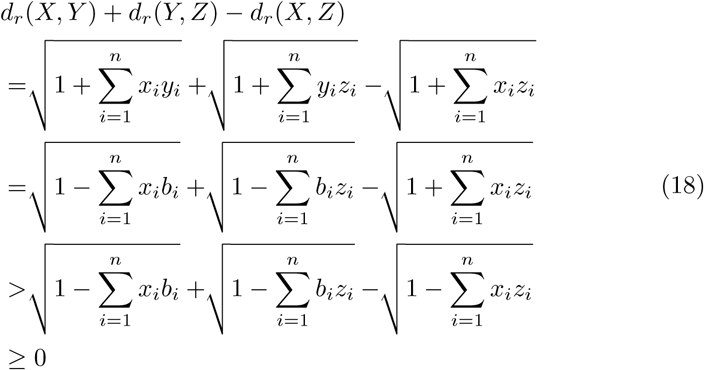

by (7)

**Case VI** When *ρ*_*XY*_ *<* 0, *ρ*_*Y*__*Z*_ *≥* 0, *ρ*_*XZ*_ *≥* 0,

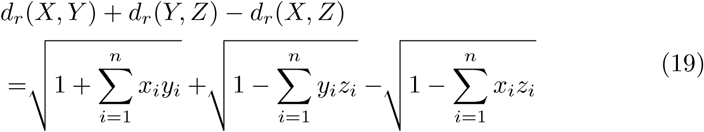

Take *a*_*i*_= −*x*_*i*_,

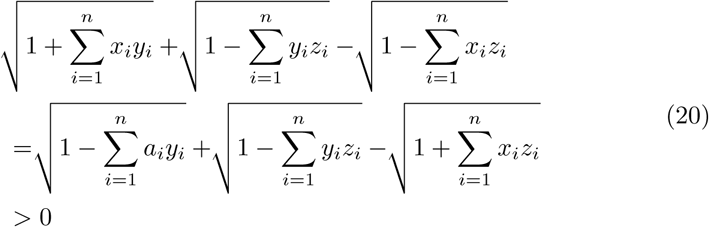

by (18)

**Case VII** The case when *ρ*_*XY*_ *<* 0, *ρ*_*Y*__*Z*_ *≥* 0, *ρ*_*XZ*_ *≥* 0, is equivalent to case VI.

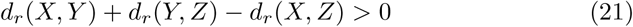

still holds.

**Case VIII** When *ρ*_*XY*_ *≥* 0, *ρ*_*Y*__*Z*_ *≥* 0, *ρ*_*XZ*_ *<* 0,

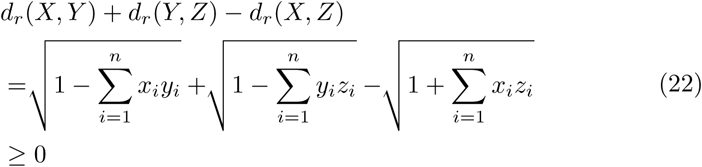

by (18)

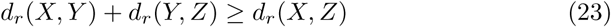

holds for any *X, Y* and *Z*.

### Proof of *d*_*r*_ fulfilling the triangular inequality for Spearman correlation as *ρ*

We define the distance of *X* and *Y* by 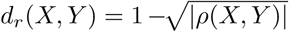, where *ρ* is the Spearman correlation.

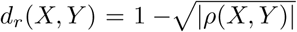, when *ρ* is the Spearman correlation, can be regarded as a special case of 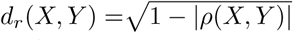, when *ρ* is Pearson correlation, where X=(*x*_1_, *x*_2_, *…, x*_*n*_), Y=(*y*_1_, *y*_2_, *…, y*_*n*_) and *x*_1_, *x*_2_, *…, x*_*n*_, *y*_1_, *y*_2_, *…, y*_*n*_ are integers. Since the inequality holds for the case of Pearson correlation, the inequality holds here.

### Proof of *d*_*r*_ fulfilling the triangular inequality for uncentered Pearson correlation as *ρ*

We define the distance of *X* and *Y* by 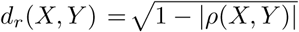, where *ρ* is the uncentered Pearson correlation coefficient.

Take *X* = (*x*_1_, *x*_2_, *…, x*_*n*_), *Y* = (*y*_1_, *y*_2_, *…, y*_*n*_) and *Z* = (*z*_1_, *z*_2_, *…, z*_*n*_). Then the triangular inequality

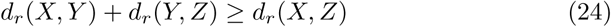

For any data from a sample, the uncentered Pearson correlation coefficient can be calculated as follows

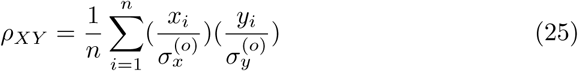

where 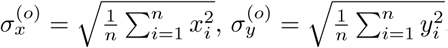.

*ρ*_*XY*_ can be written as cosine similarity,

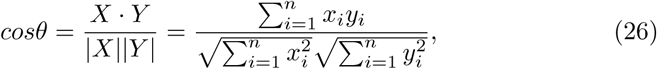

where *θ* is the angle between *X* and *Y*. Suppose *X, Y*, and *Z* are on the same plane. Let *α* denote the angle between *X* and *Y, β* denote the angle between *Y* and *Z*, such that the angle between *X* and *Z* is *α* + *β*. To prove the triangular inequality of *Γ*, we divide this into multiple cases according to the range of *α* and *β* (sign of *cosα* and *cosβ*). Suppose 0 *≤ α ≤ π*, 0 *≤ β ≤ π*,

**Case I** 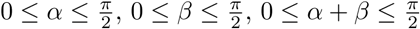,

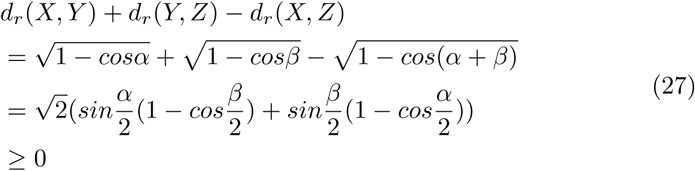

by 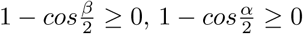.

**Case II** 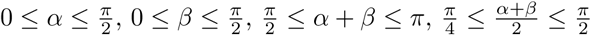

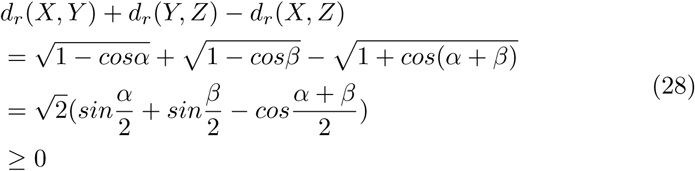

can be written as

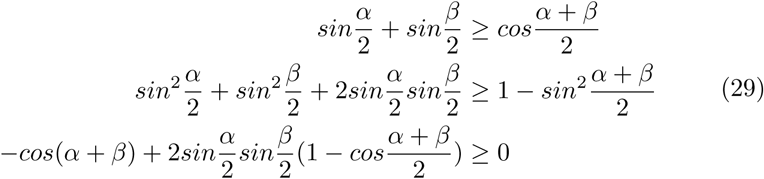

holds for 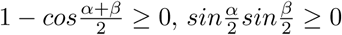 and *cos*(*α* + *β*) *≤* 0

**Case III** 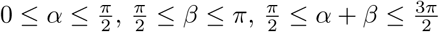

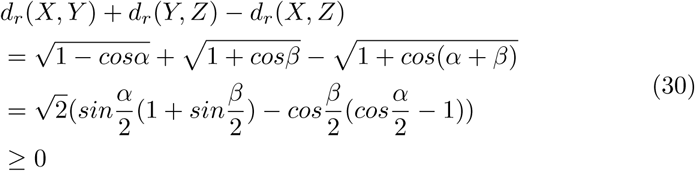

by 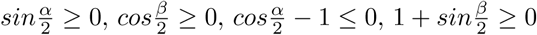

**Case IV** 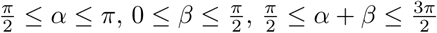 and 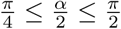

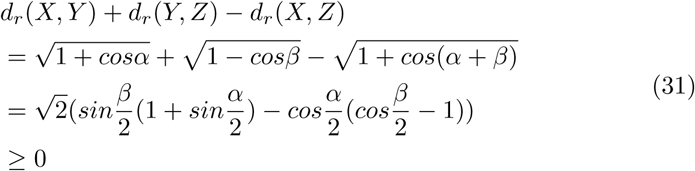

for 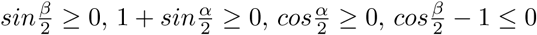.

**Case V** 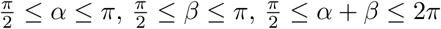 and *cos*(*α* + *β*) *>* 0 and 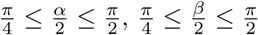

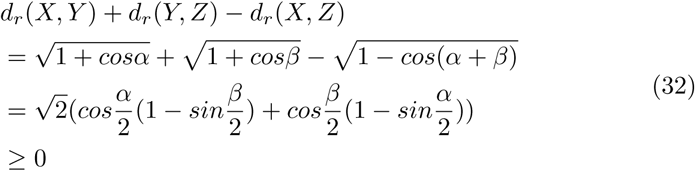

by 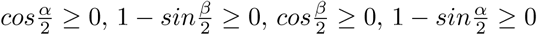

**Case VI** 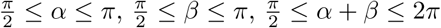 and *cos*(*α* + *β*) *<* 0, and 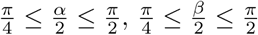

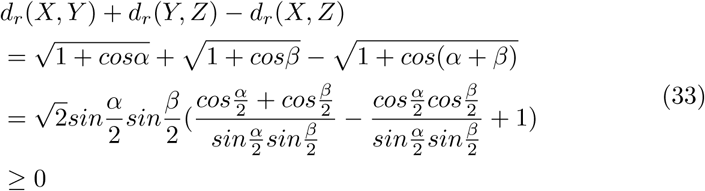

for 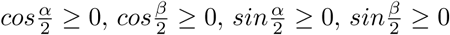

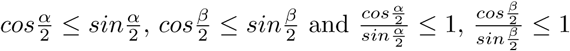

